# Integrating Functional Transdiagnostic Dimensions of Psychopathology with Cortical Organization

**DOI:** 10.64898/2026.02.02.702560

**Authors:** Alicja Monaghan, Bratislav Misic, Golia Shafiei, Kamen A. Tsvetanov, Duncan E. Astle, Richard A.I. Bethlehem

## Abstract

Childhood and adolescence represent a critical period in which neurodevelopmental psychiatric conditions emerge; traditional case-control approaches often underestimate the complexity and co-occurrence of psychiatric conditions, calling for a transdiagnostic approach as a complementary measure. Open-access data sharing initiatives provide an opportunity to decipher structural- and functional-based organisational constraints on the relationship between brain connectivity and psychopathology. Using a highly heterogenous and comorbid neurodevelopmental sample of children and adolescents aged between 6 and 17 years old at-risk of neurodevelopmental conditions (N = 174, 114 males, mean age 10.72 ± 2.21 years), and age-matched neurotypical controls (N = 27, 12 males, mean age 10.65 ± 2.07 years), we identified a multivariate association, or *latent variable*, between resting-state functional connectivity and psychopathology. We extensively benchmarked this latent construct to develop a more parsimonious account of childhood psychopathology though an analytical framework spanning biological maps, brain connectivity, and behaviour. Participant-level expression of this latent brain-behaviour association differed by diagnostic burden and symptomatology, and pre-empted longitudinal psychopathology. Whilst diagnostic status was useful for interpretation, the latent construct transcended traditional diagnostic borders, revealing a neurotypical-neurodivergent continuum. The relationship between functional connectivity and neurodevelopmental psychopathology was circumscribed by functional connectivity networks (visual, fronto-parietal, and dorsal attention) and cytoarchitectonic class (primary/secondary sensory and primary sensory cortices). The latent variable aligned with magnetoencephalography-defined electrophysiological alpha (α), high-gamma (γ), and theta (θ) frequency bands, and was enriched across cortical distributions of astrocytes and excitatory neurones. The connectivity signature was significantly aligned with the archetypal sensorimotor-to-association axis and validated in an independent sample of pre-adolescents (N = 3504), with the strongest alignment with the principal component of gene expression and myelination and were relatively less enriched in cortical regions related to language, indicating a protective effect, and more positively enriched in regions related to executive functioning, conferring greater risk of psychopathology. Together, our findings suggest that the predictive link between functional connectivity and common symptoms of neurodevelopmental psychopathology are circumscribed by underlying macroscale anatomical, functional, and cognitive-related hierarchies.

## Introduction

Multiple studies have linked different psychiatric and neurodevelopmental conditions to alterations in cortical connectivity (Buch et al., 2023; de Lange et al., 2019; Hansen et al., 2022), but how do we reconcile these findings with underlying biological mechanisms? A critical step for developing reliable biomarkers of mental health conditions is bridging multiple levels of analysis in neuroscience. One way of doing this is through harnessing recent open-access initiatives which have mapped the biological principles of cortical development (Hansen & Misic, 2025; Misic, 2025), and their links to psychopathology (Bazinet et al., 2023; Markello et al., 2022; Sydnor et al., 2021). Childhood and adolescence present a critical neurodevelopmental period for psychopathology: namely, by 14 years old, a third of youth are diagnosed with a mental health condition, increasing to almost 50% by age 18 (Solmi et al., 2022). Neurodevelopmental conditions, including autism spectrum and attention-deficit/hyperactivity conditions, are amongst the first to emerge during this critical period (Solmi et al., 2022), and feature a steady course, neurocognitive dysfunction, and multi-faceted aetiology (Thapar et al., 2017). Whilst neurodevelopmental conditions fall under the umbrella of psychopathology, in part due to their externalizing nature, they also are distinct from other conditions, including mood-based disorders, with roots in internalizing patterns (Holmes et al., 2021). The existing dogma towards conceptualising psychopathology thus far has been the diagnostic approach; wherein mental health conditions are conceptualised as homogenous and dissociable groups of separate symptoms. However, this ignores the notion that symptoms exist across a dimensional symptom space, alongside suffering from rampant comorbidity, within-condition heterogeneity, and poor discrimination between supposedly different conditions (Dalgleish et al., 2020; Marquand et al., 2016). These may be alleviated by using a *transdiagnostic* approach, which examines common symptoms across diagnostic groups, and one which already has been applied within neurodegenerative conditions such as dementia (Murley et al., 2020). Such an approach is embodied within the Research Domain Criteria (RDoC, Insel et al., 2010), which conceptualises mental health conditions as disorders of brain circuitry from which two levels of analysis extend as part of a multi-layer framework: upwards, towards phenotypic variation in psychopathology, and downwards, towards genetic, molecular, and cellular pathways.

Through a transdiagnostic lens, psychopathology may be modelled as a hierarchical taxonomy (Caspi et al., 2014; Kotov et al., 2017). At the apex is a *p* factor or general liability towards psychopathology supported by increasingly specialised strata extending from broad internalising and externalizing behaviours. One of these specialised sub-strata are neurodevelopmental conditions, thought to emerge from broad externalizing behaviours (Holmes et al., 2021). Despite the interconnected nature of sub-strata within this framework, highlighting shared variance across conditions, group-level case-control designs have been popular within the diagnostic framework. Group-level case-control designs, whilst simple to interpret, fail to reflect that patient experiences do not always map neatly onto diagnostic boundaries. Nonetheless, group-level case-control structural connectome mapping designs have demonstrated common regions are affected across neurological, psychiatric, and neurodevelopmental disorders (de Lange et al., 2019; Hansen et al., 2021). Convergence is particularly high in densely-connected ‘rich-clubs’ facilitating network communication and integration, suggesting an underlying set of core regions and systems are associated with general psychopathology (de Lange et al., 2019; Hansen et al., 2021). Such work also highlights the importance of considering brain connectivity as a *network* (reviewed by Bullmore & Sporns, 2009): abnormalities within single isolated regions are unlikely to explain psychopathology across hundreds of participants. Further, exploring cortex-wide patterns of network connectivity linked to broad psychopathology also allows for the modelling of a core concept in developmental psychopathology: *equifinality* (Cicchetti & Rogosch, 1996), which describes how multiple developmental pathways may lead to the same phenotypic manifestation. Multivariate techniques relating whole-brain connectivity and psychopathology are thus poised to capture both common and unique neurodevelopmental pathways.

Brain network analyses from resting-state functional magnetic resonance imaging (rsfMRI) have revealed multivariate links between functional connectivity networks and childhood psychopathology (Royer et al., 2024; Xia et al., 2018; Xiao et al., 2024), particularly in large-scale community-based youth cohorts, such as the Adolescent Brain Cognitive Development (ABCD) study (Casey et al., 2018) and Philadelphia Neurodevelopmental Cohort (PNC) (Satterthwaite et al., 2016). The conclusions of these studies differ in terms of analytical technique, dimensions of psychopathology captured, and sample characteristics. For example, within the PNC, loss of segregation or separation between the default mode network and executive functional networks were related to mood, psychosis, fear, and externalizing behaviour (Xia et al., 2018). In ABCD, the relationship between functional connectivity and psychopathology were patterned across distinct developmental axes. For example, connectivity within functional intrinsic connectivity networks predicted a general *p* factor of psychopathology and neurodevelopmental dimension along a sensorimotor-association axis (Royer et al., 2024). In further support of cortical axes being a guiding principle for the relationship between brain connectivity and psychopathology, in the same sample, functional connectivity along an alternative axis, the anterior-posterior axis, predicted general psychopathology and cognition (Xiao et al., 2024). The large sample sizes, and consequently higher statistical power, conferred from these consortia offers one path towards establishing more reliable brain-behaviour relationships (Marek et al., 2022). However, a complementary path are focused samples which oversample from clinically-relevant populations, and thus maximise signal-to-noise ratios (Gratton et al., 2022). One example is the Centre for Attention, Learning, and Memory (Holmes et al., 2019) dataset: a transdiagnostic, developmental sample spanning childhood and adolescence, recruited with single, multiple, or no formal diagnoses, but on the basis of broad difficulties across attention, learning, or memory.

Whilst psychopathology itself is heterogenous, existing work examining the relationship between functional connectivity and psychopathology conceptualises functional activation within and between brain regions as a homogenous node. However, an additional layer of complexity exists beyond the presence of functional activation, namely biological embeddings and context (see Bazinet et al., 2023). Each functional node is embedded within discrete and continuous hierarchical organisational systems. Discrete classifications include those based on cytoarchitectural similarity (von Economo & Koskinas, 1925), laminar differentiation (Mesulam, 2000), and intrinsic resting-state functional connectivity networks (Yeo et al., 2011). Continuous gradients of organisation spanning microstructure, macrostructure, functional connectivity, and evolutionary constraints were recently formalised as a sensorimotor-association axis (S-A) (Sydnor et al., 2021). This axis differentiates between lower-order, unimodal, highly-myelinated sensorimotor cortices supporting externally-directed behaviour, and the higher-order, transmodal, lightly-myelinated association cortices supporting internally-directed thought and cognition. Evaluating alignment between a feature of interest and each composite map of the S-A axis provides useful clues as to which developmental processes and structures may be implicated, or not, in the emergence or maintenance of this feature. Crucially, each organisational feature, such as correlated gene expression and canonical magnetoencephalography frequency bands, is non-redundant (Hansen et al., 2023), meaning that whilst certain organisational patterns are shared, each layer provides a unique perspective. Therefore, contextualising the relationship between FC and psychopathology using multi-modal cortical organisational principles may reveal specific biological patterns linked to risk of or resilience against psychopathology.

In the current study, we explored the multivariate relationship between functional connectivity and neurodevelopmental psychopathology in a transdiagnostic, developmental cohort (N = 201, 6-17 years old) from the Centre for Attention, Learning and Memory (CALM; Holmes et al., 2019). A key feature of CALM, and in contrast to typically developing cohorts drawn from the general population, most children were at-risk of neurodevelopmental conditions but undiagnosed. Consistent with RDoC recommendations (Insel et al., 2010), and utilising open-access initiatives providing cortical biological maps (Markello et al., 2022; Shafiei et al., 2023; Sydnor et al., 2021), we implemented a multi-level analytical framework spanning biology, neural circuitry, and behaviour. Using extensive cross-validation and reliability analyses in a highly heterogenous sample, and benchmarking FC-psychopathology relationships with cortical biological maps using conservative spatial permutation tests (Alexander-Bloch et al., 2018; Váša & Mišić, 2022), and with broader cognition and behaviour using external meta-analytic engines (Yarkoni et al., 2011), we aimed to delineate which functional and biological systems are most at-risk or resilient to psychopathology. In keeping with the overlapping cortical organisational principles described previously, such as the somatosensory-association axis (Sydnor et al., 2021), we hypothesized that the relationship between FC and psychopathology would be differentially constrained by underlying structural hierarchies. In other words, we predicted that the relationship between FC and psychopathology would be non-random, rendering certain, particularly higher-order associative, systems more vulnerable than others. Through embracing the complexity of psychopathology phenotypes, using a sample more representative of real-life practice, and leveraging open-access initiatives providing high-resolution biological maps of the cortex, we hope to begin to reconcile alterations in high-resolution cortical connectivity in neurodevelopmental conditions with their biological underpinnings.

## Methods

### Participants

957 children aged between 5 and 18 years old were recruited from schools, primarily within the South East of the United Kingdom, with ethical approval from the National Health Service local authority (reference 13/EE/0157), following a prior published protocol (Holmes et al., 2019). Children were included if they displayed difficulties within attention, learning, and/or memory, indicating a general risk towards neurodevelopmental conditions, regardless of diagnosis or comorbidities. In addition, children must be native English speakers without a known genetic condition affecting cognition or uncorrected sensory problems. Within this larger behavioural subset, 348 children underwent resting-state fMRI scans, from which we obtained low-motion processed data for 201. Full recruitment and processing details are available elsewhere (Holmes et al., 2019; Jones et al., 2021). In brief, the 201 participants were aged between 6 and 17 years old (*M* = 10.71 years, *SD* = 2.19 years), with 63% males, and an average in-scanner motion of 0.21 mm (*SD* = 0.09mm). From the 107 participants with ethnicity data, most were White (88.76%), with 10.28% having multiple ethnicities and 0.93% Black. Of the 174 participants at risk of neurodevelopmental conditions, most were referred due to poor scholastic performance (31.21%), followed by issues in attention (29.48%), memory (15.03%), literacy (12.14%), language (9.25%), and mathematics (2.89%). 67.82% were referred by education specialists, 27.01% were referred from paediatric or adolescent mental health services, and the remainder were referred by speech and language therapists. Of the 174 at-risk participants, most did not have a formal diagnosis (62.64%). Of those who did, most were neuro-developmental, including ADHD (18.39%), dyslexia (9.77%), and autism (7.47%), with 11.94% having two comorbidities or more. The remainder (N = 27) formed the age-matched comparison group and were recruited from the same schools as the at-risk children, but without a referral to CALM. None of the control participants, bar one, had a formal diagnosis.

The at-risk and control participants were well matched on demographic characteristics. For example, a 2 × 2 chi-square test of sample (2 levels – referred or control) and sex (2 levels – male or female) showed no significant relationship between sample and sex [χ^2^(1, *N* = 201) = 0.62, *p* = 0.43]. A 5 × 2 chi-square test of sample and ethnicity (3 levels – White, Black, or multiple ethnicities) showed no significant relationship between sample and ethnicity [χ^2^(2, *N* = 201) = 5.02, *p* = 0.08]. We compared continuous measures between samples using a series of independent sample t-tests. The at-risk and control participants did not differ in terms of deprivation [*t*(199) = 1.83, *p* = 0.07)], as measured by the postcode-based index of multiple deprivation. However, there was a significant difference between samples in terms of in-scanner head motion [*t*(199) = 2.07, *p* = 0.04, Cohen’s *d* = 0.43], as measured by mean framewise displacement, with the at-risk participants [*M* = 0.21mm, *SD* = 0.10mm] having greater motion than the comparison group [*M* = 0.17mm, *SD* = 0.08mm]. Further, there was a significant difference between samples in terms of fluid reasoning [*t*(199) = - 4.37, *p* = 2.01 × 10^-5^, Cohen’s *d* = -0.90], as measured by the matrix reasoning subscale of the Wechsler Abbreviated Scale of Intelligence (Wechsler, 2011), whereby the at-risk participants exhibited lower fluid reasoning [*M* = -0.58, *SD* = 1.03] than their neurotypical counterparts [*M* = 0.36, *SD* = 1.01].

### Functional data acquisition, pre-processing and connectome construction

We used T1w and resting-state FC data in the current study, both of which were obtained from a 3-Tesla Siemens Prisma MRI scanner, with 32-channel quadrature head coil, at the MRC Cognition and Brain Sciences Unit (University of Cambridge), during a single session, following the protocol described prior (Holmes et al., 2019). Participants were instructed to stay awake and close their eyes. 270 volumes of T2*-weighted fMRI data were acquired using a gradient-echo planar imaging sequence, with each volume containing 32 axial slices, with repetition time (TR) of 2000 milliseconds, echo time (TE) of 30 milliseconds, 3mm isotropic voxel resolution, across 9 minutes and 6 seconds. These were co-registered to 192 slices of T1-weighted images collected using a magnetization prepared rapid gradient echo imaging sequence, with TR of 2550 milliseconds, TE of 3.02 milliseconds, 1mm isotropic voxel resolution, across 4 minutes and 32 seconds. Through prior work (Jones et al., 2021), we obtained fMRI data processed using fMRIprep version 1.5.0 (Esteban et al., 2019), the full pipeline details of which are available at https://fmriprep.readthedocs.io/en/latest/workflows.html. In brief, this implemented slice-timing correction, rigid-body realignment, T1w skull-stripping and spatial normalization, alignment of the functional image to the T1w reference using boundary-based co-registration, and normalisation to MNI template space. Head motion was corrected for through motion spike regression, anatomical component correction, and regression of 24 head motion parameters (Jones et al., 2021). The pre-processed time-series were parcellated to the 400-region Schaefer atlas (Schaefer et al., 2018), and the functional connectome quantified as the Pearson correlation coefficient between pairwise regional timeseries. As a sensitivity analyses, we also obtained identically-processed connectomes in the 100-node Schaefer parcellation, from a prior study (Jones et al., 2022) in which the edge weights had undergone a Fisher’s z transformation.

### Conners psychopathology scale

Participants’ parents completed the short-form version of the 3^rd^ Edition of the Conners scale (Conners et al., 2011), in which they used a 4-point Likert scale to report how much they believed each statement to be true of their child in the past month, ranging from not true at all (0 points) to very much true (3 points), across 31 items examining executive functioning, learning difficulties, peer relations, aggression, inattention and hyperactivity. An additional 12 items measured positive and negative reporting bias (6 each): scores of 5 or above indicated an overly positive or negative response bias, respectively. The Conners scale particularly targets symptoms related to Attention Deficit Hyperactivity Disorder (ADHD) and comorbid conditions, including conduct disorder.

### Brain-behaviour multivariate partial least squares (PLS) analysis

Using the pyls package in Python (https://github.com/rmarkello/pyls), we implemented PLS to relate edge-wise FC with psychopathology items. PLS is a multivariate statistical technique which uses singular value decomposition (SVD; Eckart & Young, 1936) to produce orthogonal latent variables which maximises the input-output covariance. As shown in **Equation 1**, Χ represents the participant-level edge-wise FC (79,800 × 201), whilst Υ represents the participant-level psychopathology items (31 × 201).

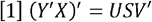

SVD on the covariance matrix Υ′Χ produces 3 outputs. The first two are the singular vectors U and V, the *i*-th columns of which represent the latent variable *i*. The contribution of individual functional connections (FC-LV) and psychopathology items (psychopathology-LV) to the latent variable are represented by U and V. Positive weights suggest a positive FC-psychopathology relationship, conferring risk, whilst negative weights suggest a negative FC-psychopathology relationship, conferring resilience. The third output is the singular values, the *i*-th diagonal of which are proportional to the covariance captured by the *i*-th latent variable. The covariance η explained by each latent variable *i* is equal to the squared singular value σ corresponding to that variable, divided by the sum of all squared singular values, where j is the total number of singular values (see **Equation 2**).

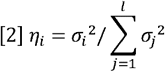

Brain and behavioural subject scores are quantified for each participant by projecting left and right singular vectors back onto the original data. These scores represent the extent to which the participant expresses the FC pattern or psychopathology behavioural pattern of the LV, respectively. Following prior work (Mišić et al., 2016), we used bootstrapping to assess the reliability of the brain weights. The rows of the FC and psychopathology matrices, corresponding to participants, were randomly subsampled with replacement, and SVD repeated to produce new brain weights. This was repeated 5000 times to create a distribution of brain weights. The empirical brain weights were then normalised by the standard error of the bootstrapped distribution to create a bootstrap ratio (BSR), such that functional connections with a large BSR were both large in magnitude and small in variability across participants (as in Mišić et al., 2016). This is an alternative to thresholding the participant-level FC matrices before PLS, which may otherwise mask weak functional connections that are consistently related to psychopathology.

To assess the statistical significance of each latent variable, we utilised a permutation testing procedure repeated 5000 times. For each permutation, the rows of the FC matrix were shuffled, thereby breaking the FC-psychopathology relationship, and the SVD recalculated. The one-tailed permuted p-value was calculated as the proportion of times the permuted singular value exceeded the empirical singular value. To assess the out-of-sample generalisability, we implemented 5-fold cross-validation stratified on age (2 bins, split at 11.5 years old) and sample (at-risk or comparison), comprising 4 stratification groups, repeated across 1000 iterations. For each iteration, the connectivity and psychopathology matrices were split into 5 folds, using the two stratification variables. 80% of the data was assigned to training, and the remainder to testing. The rows of the psychopathology matrix were shuffled to produce a null distribution. The PLS was conducted on the training data. For each latent variable, iteration, and fold, the × and Y subject scores from the empirical PLS were correlated. The training weights were then projected to the test data for the connectivity and psychopathology matrices separately and correlated. The same procedure was applied for the PLS trained on the null distribution. Once averaged across all folds for each latent variable, this formed the mean training, test, and null correlations, respectively. The cross-validated one-tailed p-value was the proportion of times that the null distribution correlation exceeded that derived from the empirical testing distribution.

To assess the stability of the first latent variable linking FC and psychopathology, we implemented a split-half reliability procedure (Kovacevic et al., 2013). Specifically, for each of 1000 iterations, one participant was randomly removed to create an even sample of 200 participants. Using the same stratification criteria as above, the participants were stratified into two equal samples, and the cross-covariance matrix calculated for each, comprising *R*_*1*_ and *R*_*2*_ respectively, and SVD computed to derive connectivity (*U*_*1*_ and *U*_*2*_) and psychopathology (*V*_*1*_ and *V*_*2*_) singular vectors for each half. The stability of the connectivity singular values was the mean *U*_*1*_-*U*_*2*_ correlation across iterations, and vice versa for the *V*_*1*_-*V*_*2*_ correlations representing the psychopathology signature.

### Alignment with the S-A axis

To examine the distribution of brain BSRs, we used the sensorimotor-association (S-A) axis derived from the distribution of 10 evolutionary, functional, microscale, and macroscale cortical biological maps (Sydnor et al., 2021). Relative to the sensorimotor pole, the association pole exhibits less myelination, measured by a T1-weighted to T2-weighted ratio from adult normative structural data (Glasser & Van Essen, 2011); greater involvement in unimodal, lower-order processing, measured by the principal gradient of adult normative functional data (Margulies et al., 2016); and greater macaque-to-human evolutionary expansion (Hill et al., 2010). Further, the association pole features greater areal scaling relative to the whole brain (Reardon et al., 2018); lower glucose metabolism, measured by positron emission tomography (Vaishnavi et al., 2010); greater cerebral blood flow, measured by arterial spin labelling (Satterthwaite et al., 2014); and anchoring one end of a principal gradient of cortical gene expression variation, based on micro-array data from the Allen Human Bran Atlas (Burt et al., 2018; Hawrylycz et al., 2012). The final features of the association pole within the S-A axis includes greater involvement in internal cognitive and behavioural processes, such as thought and emotion, measured by the principal component of Neurosynth meta-analytic functional activations (Yarkoni et al., 2011); greater proportion of internopyramidisation or infragranular feedback connectivity, measured by BigBrain histology (Amunts et al., 2013; Paquola et al., 2020); and greater cortical thickness.

### Independent validation of the connectivity signature in ABCD

Following the same procedure as Voldsbekk and colleagues (2023), we formally tested the replicability of the identified S-A axis alignment with that of an independent ABCD study linking general psychopathology in 3,504 participants with multimodal neuroimaging, including resting-state FC (Royer et al., 2024). Specifically, Royer and colleagues (2024) conducted principal component analysis (PCA) on resting-state FC, which was then subjected to a PLS. Consequently, rsFC PCA coefficients and rsFC loadings were made available, and are also provided in our GitHub repository. We restricted our comparison to 400 cortical nodes, excluding the 19 subcortical nodes included in the independent study (Royer et al., 2024). First, we reshaped the provided ABCD 400 × 400 × 256 component PCA weights into a vector of size 160,000 × 256. This was multiplied by the raw CALM functional connectomes with the same number of edges but with 201 participants, producing a vector of size 201 × 256. For interpretability, and as PLS-derived signs are arbitrary, we flipped the ABCD PLS vector. We then multiplied this by the ABCD rsFC PLS loadings to create a connectivity vector of ABCD-derived connectivity in CALM. This was not significantly correlated with our PLS-derived psychopathology participant-level scores, potentially due to different measures used, suggesting replicability of the connectivity signature only.

### Null models

Spatial autocorrelation is a core feature of brain organisation, where spatially proximal regions are more similar that spatially distant regions, above chance. Spin tests are a form of null model which assess whether the spatial correspondence between two brain maps exceeds that expected by spatial autocorrelation (Markello & Misic, 2021). In the spatial permutation framework (Alexander-Bloch et al., 2018; Váša et al., 2018), the brain map is projected onto a sphere whose coordinates are randomly rotated 5000 times, projected back onto the anatomical surface, and each region reassigned once according to Euclidean distance. Thus, the original feature values and spatial autocorrelation are preserved, but the underlying relationship with cortical geometry is otherwise disturbed. We used these methods for all parcellated dense matrices. The only exception was for the genetic gradients derived from the Allen Human Brain Atlas (AHBA), parcellated into 400 nodes, using custom code from a prior study (Dear et al., 2024). Restricted to the left hemisphere, 44 out of the 200 regions did not have genetic data. To preserve this sparsity, we used the Cornblath null model (Cornblath et al., 2020). This projects the parcellated data up to the *fsaverage* surface (41k vertices), spins the data on the sphere, and re-averages back to the original parcellation.

### Cell-type gene expression, cytoarchitecture, laminar differentiation and Neurosynth functional activations

We obtained digitized maps spanning the 7 von Economo and Koskinas cytoarchitectonic differentiation classes, in the Schaefer 400-node parcellation (Hansen et al., 2024; Vértes et al., 2016; von Economo & Koskinas, 1925). Note, these were extended (Vértes et al., 2016) from the original 5 classes to include the limbic cortex (comprised of entorhinal, retrosplenial, presubicular, and cingulate cortex) and insular cortex (comprised of granular, agranular, and dysgranular regions). Digitized maps spanning the 4 levels of laminar differentiation were also obtained in the 400-node resolution, spanning unimodal, paralimbic, heteromodal association, and idiotypic cortices (Mesulam, 1998). We retrieved cell type enrichment maps for 7 canonical cell types, calculated as the average expression of cell-type specific gene sets derived from cell type deconvolution in the AHBA (Seidlitz et al., 2020; Shafiei et al., 2023). Adult normative magnetoencephalography (MEG) maps, parsed into 6 canonical frequencies and intrinsic timescale, were obtained from the S1200 release of the Human Connectome Project (Van Essen et al., 2013), and parcellated from the *fsaverage* surface (4k density) using *neuromaps* (Markello et al., 2022).

Neurosynth is a meta-analytic engine which uses text-mining to compile voxel activation coordinates for fMRI data associated with specific cognitive and neural states (Yarkoni et al., 2011), across almost 15,000 studies. The result is a series of spatial association maps denoting the probability that a specific high-frequency term appeared co-occurred with functional activation in that region. Such probabilistic associations do not incorporate directionality (Hansen et al., 2021). To narrow our focus to cognition and limit the number of statistical comparisons, we examined the relationship between the PLS-derived cortical bootstrap ratios and 123 functional activation term maps from the Cognitive Atlas, grouped into 11 categories (Poldrack et al., 2011).

### Three components of cortical gene expression

To explore how the relationship between FC and psychopathology may extend beyond the sensorimotor-association axis, we derived three cortical gene expression gradients following prior work (Dear et al., 2024), using code adapted for the Schaefer 400-node parcellation. In brief, we used the *abagen* toolbox (Markello et al., 2021) in Python to create a gene expression matrix for the left hemisphere from probes present in at least 3 out of 6 donors, retaining genes with differential stability of at least 50%, followed by diffusion-map embedding to derive 3 components. Full methodological details are provided elsewhere (Dear et al., 2024). Note that due to the different processing parameters for the principal component of gene expression forming the S-A axis and that derived in this case differed slightly, the principal component from both pipelines is strongly correlated [*r* = 0.949, *p* = 3.542 × 10^-79^ ] but not identical.

## Results

### The multivariate pattern capturing covariance between FC and psychopathology is organised non-randomly

We related item-level psychopathology with edgewise resting-state functional connectivity using partial least squares, which seeks to maximise the covariance between predictors and responses (**Figure 1a**). One latent variable (LV) accounted for 43% of the psychopathology-FC covariance (permuted *p* = 0.012), with statistical significance determined by comparing the empirical singular value with that of 5,000 permutations (**Figure 1b**). Whilst the second LV was also statistically significant (permuted *p* = 0.022), capturing 11.63% of covariance, we focus on the first LV henceforth as it accounted for more than four-fold covariance. Since the first LV captured most covariance, this suggests that we have captured a broad neurodevelopmental factor of psychopathology. Projecting the PLS-derived weights onto the original FC and psychopathology data produces brain and behavioural subject scores, respectively. These index to what extent each participant expresses the brain connectivity and behavioural pattern identified in the first LV. By design, these are highly correlated (Pearson’s *r*(199) = 0.646, *p* = 4.07 × 10^-25^ ) such that participants who highly express the connectivity pattern also highly express the behavioural signature (**Figure 1c**).

**Figure 1.**
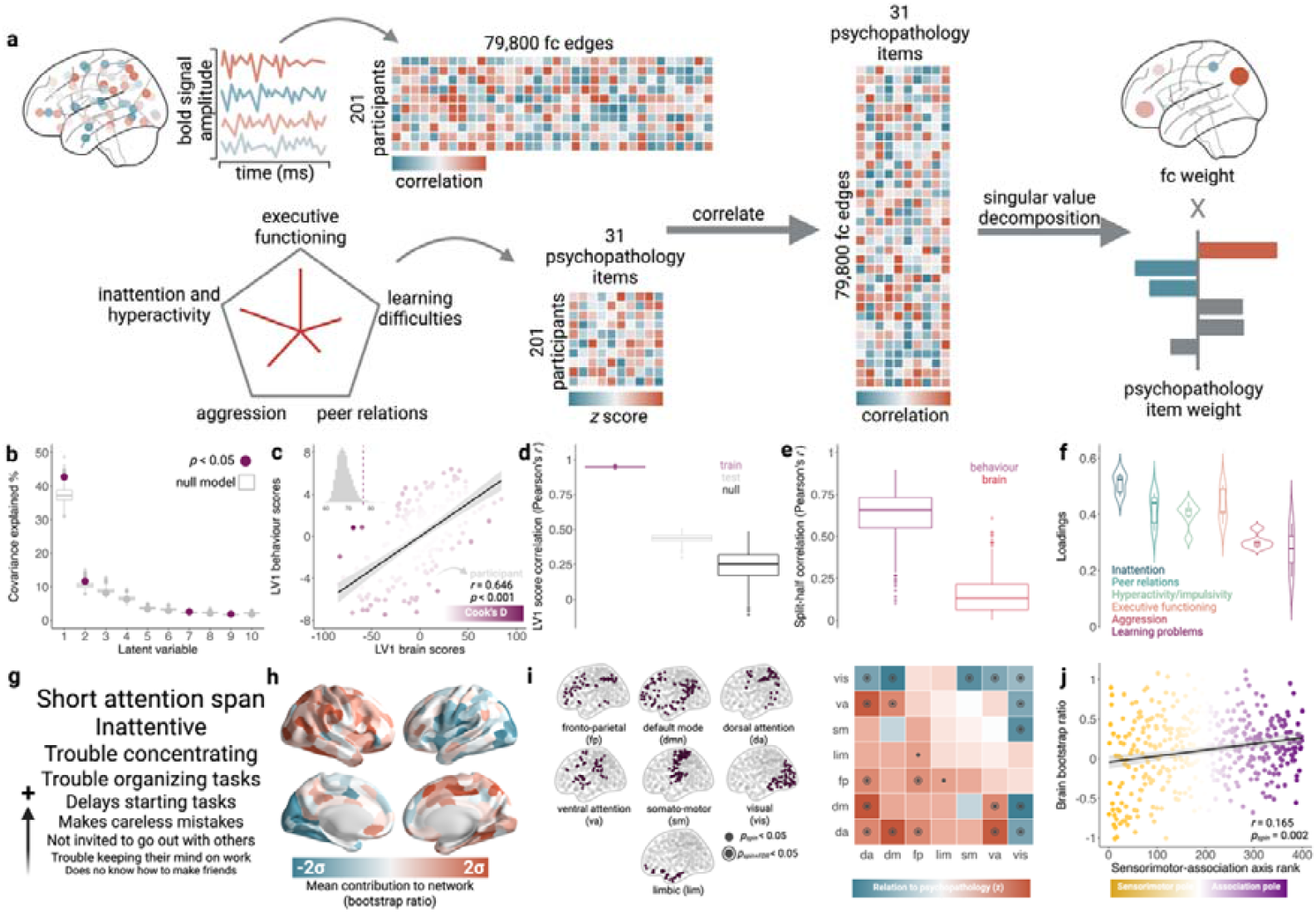
Relating functional connectivity to psychopathology. **a**. Across 201 CALM participants (174 referred, 27 controls), PLS was used to relate edgewise resting-state functional connectivity to item-level psychopathology from the Conners scale, spanning executive functioning (n = 5), learning difficulties (n = 5), peer relations (n = 5), aggression (n = 5), and inattention/hyperactivity (n = 6). The participant-level FC and psychopathology matrices were cross-correlated and subjected to singular value decomposition. This produces a series of latent variables (LV), whereby FC and psychopathology are weighted and linearly combined to maximise their covariance. **b**. The first LV accounted for 43% of FC-psychopathology covariance. Statistical significance was assessed by randomly permuting the original data to form a null model, and comparing the magnitude of the singular values derived from the null model with the empirical model. c. Brain and behavioural scores are the product of the original data projected back onto the PLS-derived weights and represents the extent to which each participant expresses the connectivity and behavioural patterns identified by the PLS. The influence of brain scores on behavioural scores was assessed using linear regression and quantified by Cook’s distance (Cook, 1977), visualised in purple. **c.** The out-of-sample predictive value of the FC-psychopathology relationship was assessed using 5-fold cross-validation, repeated 1000 times, and stratified by sample (referred or control) and age (4 equally sized bins between 6 and 17 years old). **e.** To assess the reproducibility of the first LV, we conducted split-half resampling, repeated 1000 times, whereby the entire sample was randomly split in half, the PLS repeated in each half, and the behavioural scores from each half correlated, followed by the brain scores. **f.** The distribution of statistically significant psychopathology item loadings with non-zero confidence intervals are shown as violin plots using a gaussian kernel. All items had a positive loading. **g.** Top-loading psychopathology items. Note, some were shortened for brevity. **h.** The contribution of each brain region to the identified FC-psychopathology connectivity signature is visualised on the inflated cortical surface at 95% confidence intervals, as bootstrap ratios, using the Schaefer 400-node parcellation (Schaefer et al., 2018). These are the brain weights divided by the standard error of 5000 bootstrap resamples, indexing the reliability of the contribution of each region. The original 4950 × 1 matrix was reshaped into a symmetrical 2D matrix, and bootstrap ratios averaged across regions. **i.** The z-scores, given as the empirical bootstrap ratio relative to that of a null distribution with preserved spatial autocorrelation, were converted to p-values. The false discovery rate (FDR) was corrected across the upper triangle of the matrix, including the diagonal, comprising 28 comparisons. The full matrix is shown for visualisation only. **j.** The distribution of nodal functional connectivity contributions to psychopathology follows a sensorimotor-to-association axis (Sydnor et al., 2021), the significance of which was assessed by comparing the empirical correlation with null correlations from 5000 null models in which spatial autocorrelation was preserved. Individual points represent brain regions and are coloured by their sensorimotor-association axis rank.

We conducted several analyses to assess the robustness, reliability, and stability of the FC-psychopathology relationship. First, to assess whether any individual participants have an unduly large influence on the slope of the FC-psychopathology relationship, we calculated Cook’s distance (Jones et al., 2022) after regressing behavioural scores onto brain scores. This is a leave-one-out measure which examines the change in slope when each individual participant is removed. These values were particularly small (maximum distance = 0.051), substantially below the threshold of 1 above which a data point is considered influential (Hair Jr. et al., 2018). We checked this separately in children flagged for potential negative reporting bias (N = 17, maximum distance = 0.009) and for those receiving ADHD medication (N = 22, maximum distance = 0.019), both of which displayed small distances. We therefore concluded that the FC-psychopathology relationship identified in the PLS was not driven by any individual children (or subset) but rather reflects an association across the entire sample. Second, we assessed the out-of-sample brain-behaviour relationships using 5-fold cross-validation, stratified by sample (control or referred) and age (two equally sized bins between 6 and 17 years old). For each split, we trained the PLS using 80% of the data, and projected these weights back onto the remaining 20%, repeated 1000 times. The distribution of training, test, and null correlations are shown in **Figure 1d**. The mean out-of-sample Pearson’s correlation was *r*(998) = 0.186 and was both significant against a permuted null model (*p* = 0.001) and significantly greater than 0 (*t*(999) = 195.356, *p* < 0.001). Third, we estimated the stability of the FC-psychopathology relationship, we conducted split-half resampling. In this framework, the data is randomly split into halves and the PLS repeated in each half. The brain scores from each half are correlated, and likewise for behavioural scores. The distribution of correlations across 1000 splits are shown in **Figure 1e**. In line with prior brain-behaviour studies which have examined relationships between mood and FC (Mirchi et al., 2019), alongside structural connectivity and schizophrenia (Kirschner et al., 2020), we demonstrated higher split-half reliability for behavioural scores (mean *r* = 0.628 [95% confidence interval (CI): 0.619, 0.637]) compared to connectivity (mean *r* = 0.149, 95% CI: 0.142, 0.156). This likely reflects the behavioural data having fewer variables and less noise than the connectivity data. Whilst such split-half reliability is relatively modest (Helmer et al., 2024), to which we return in the **Discussion**, subsequent independent validation of the connectivity signature, enrichment of the latent variable with biological maps under strict permutation strategies, longitudinal prognostic value of connectivity scores for psychopathology, and replication with regularized sparse CCA suggests that the derived FC-psychopathology relationship is stable and robust to noise.

Having established the robustness of the FC-psychopathology relationship, we examined the contribution of individual psychopathology items. 29 of the 31 psychopathology items loaded significantly (non-zero confidence intervals) onto the first LV, all of which positively (**Figure 1f**). An ANOVA and post-hoc Tukey test showed a significant effect of category on loadings (*F*(5, 25) = 13.832, *p* = 1.588×10^-6^). Inattention loaded most strongly (*M* = 0.510, *SD* = 0.036), significantly more so than aggression (*diff* = 0.206, *p* = 5.71 × 10^-4^) and learning problems (*diff* = 0.311, *p* = 1.00 × 10^-6^). Learning problems overall loaded least strongly (*M* = 0.199, *SD* = 0.125), and significantly less so compared to peer relations (*diff* = -0.219, *p* = 2.64 × 10^-4^), executive functioning (*diff* = -0.236, *p* = 9.60 × 10^-5^), hyperactivity/impulsivity (*diff* = -0.201, *p* = 4.50 × 10^-4^), and inattention (*diff* = -0.311, *p* = 1.00 × 10^-6^). Of the 5 top-loading items (**Figure 1g**), 3 related to inattention, namely “has a short attention span” (*r* = 0.547, 95% CI: 0.456, 0.632), “inattentive” (*r* = 0.535, 95% CI: 0.440, 0.623), and “has trouble concentrating” (*r* = 0.524, 95% CI: 0.422, 0.615) whilst 2 related to executive functioning, namely “has trouble organizing tasks or activities” (*r* = 0.504, 95% CI: 0.385, 0.605) and “has trouble getting started on tasks or projects” (*r* = 0.490, 95% CI: 0.373, 0.592). Of the 5 lowest-loading items, 2 related to learning problems, namely “cannot grasp arithmetic” (*r* = 0.177, 95% CI: 0.036, 0.316) and “does not understand what they read” (*r* = 0.278, 95% CI: 0.149, 0.396), whilst 3 related to aggression, namely “tells lies to hurt other people” (*r* = 0.280, 95% CI: 0.154, 0.387), “starts fights with others on purpose” (*r* = 0.288, 95% CI: 0.165, 0.401), and “threatens to hurt others” (*r* = 0.299, 95% CI: 0.187, 0.403). In support of the behavioural component reflecting a general *p*-factor of psychopathology, the PLS-derived behavioural scores were strongly correlated to *p*-factor loadings derived from separate questionnaires (Holmes et al., 2021) for 67 common participants (*r* = 0.833, *p* = 2.139 × 10^-18^).

The relative magnitude and reliability of the contribution of regions to the LV connectivity pattern were quantified by nodal bootstrap ratios (**Figure 1h**), equal to the individual weight of a region divided by the bootstrap-estimated standard error. Regions in which FC conferred resilience against higher psychopathology, indexed by negative bootstrap ratios, were centred in the left hemisphere within lower-order visual and somato-motor cortices. Regions in which functional connectivity predicted higher psychopathology, indexed by positive bootstrap ratios, were centred in higher-order portions of the right hemisphere, within the posterior dorsal attention network and the medial ventral attention network. We then examined the extent to which bootstrap ratios were constrained within and between 7 functional resting-state networks (RSNs, Yeo et al., 2011). To do so, we permuted the RSN labels using a spatial null model 5000 times, recalculated the mean bootstrap ratio for each RSN-RSN pair, and expressed the empirical mean bootstrap ratio as a z-score using the mean and standard deviation of the null distribution (**Figure 1j**). Higher psychopathology was associated with increased functional connectivity within and between higher-order RSNs, such as the frontoparietal, default-mode, and dorsal attention networks, whilst lower psychopathology was associated with increased connectivity between the lower-order visual RSN and others, particularly the default-mode and somato-motor RSNs. Note that when we restricted our analysis to the mean nodal bootstrap ratios, rather than the full edge-wise connectome, we observed significant positive enrichment within the fronto-parietal [*p* = 0.012, Cohen’s *d* = 2.48] and dorsal attention RSNs [*p* = 0.010, Cohen’s *d* = 2.524] conferring risk to psychopathology, alongside significant negative enrichment within the visual RSN [*p* < 0.001, Cohen’s *d* = -3.92] conferring resilience.

### Alignment with the sensorimotor-association axis and independent validation of the connectivity signature

In line with prior work in 3,504 ABCD participants aged between 9 and 10 years old (Royer et al., 2024), and to contextualise our findings in terms of broader organisational cortical axes, we demonstrated that the nodal distributions of bootstrap ratios were significantly positively correlated with a sensorimotor-to-association axis (*r* = 0.165, *p*_*spin*_ = 0.002), the maturation of which has been associated with psychopathology (Royer et al., 2024; Sydnor et al., 2021). Regions at the association end were stronger positive predictors of psychopathology than at the sensorimotor end (**Figure 1j**). Such replicability suggests that, despite the high dimensionality of the current study, overfitting was limited. Note that, whilst the this axis is adult-derived, as outlined by Sydnor and colleagues (2021), brain development unfolds along this stable axis, such as alignment of functional connectivity across childhood and adolescence (Luo et al., 2024), and thus can be used as a theoretical and experimental scaffold upon which to interpret developmental events. As carried out by Voldsbekk and colleagues (2023), to formally test such replicability, and as detailed in the independent validation methodology sub-section, we projected the ABCD PLS rsFC loadings onto the raw CALM functional connectomes to create an ABCD-derived connectivity vector in CALM. This was significantly correlated with the original CALM brain scores (*r* = 0.358), suggesting that participants in both cohorts shared similar connectivity signatures. We assessed the statistical significance of this relationship in two ways. The first was a participant-level permutation procedure (Voldsbekk et al., 2023), which rejected the null hypothesis that there was no significant relationship between participant-level ABCD-derived connectivity scores and the original CALM connectivity scores (*p* = 0.001). The second was a FC-level permutation procedure, where surrogate functional connectomes preserving spatial autocorrelation (Váša et al., 2018) were created for each CALM participant by applying spin indices to rows and columns, after which the PLS/PCA multiplication procedure was repeated. This allowed us to reject the null hypothesis that applying PLS/PCA ABCD weights onto noise produces a connectivity vector significantly correlated with the original (*p* = 0.001).

### Exploring characteristics of brain and behavioural scores

Having scrutinised the reliability and spatial patterning of the LV linking FC and psychopathology, we then examined how participant-level expression of the LV connectome pattern (brain scores), and psychopathology pattern (behavioural scores) vary by demographics, diagnostic burden, and symptom profiles. Using a series of linear regressions (**Figure 2a**), we found that males had significantly higher brain (β = 0.281, 95% CI: [0.003, 0.559], *p* = 0.474) and behavioural (β = 0.407, 95% CI: [0.138, 0.677], *p* = 0.003) scores than females, possibly reflecting the over-representation of males in CALM. As expected, head motion significantly predicted stronger brain (β = 0.301, 95% CI: [0.169, 0.433], *p*= 1.22 × 10^-5^) and behavioural scores (β = 0.178, 95% CI: [0.050, 0.307], *p* = 0.007). In subsequent sensitivity analyses, we demonstrate that our findings are not reliant on head motion. Higher deprivation was linked to stronger behavioural scores (β = 0.294, 95% CI: [0.130, 0.458], *p* = 4.98 × 10^-4^ ), but not connectome scores. Finally, lower fluid reasoning was linked to lower brain (β = -0.125, 95% CI: [-0.250, -0.001], *p* = 0.048) and behavioural scores (β = -0.206, 95% CI: [-0.327, -0.086], *p* = 0.001). Age did not significantly predict either brain or behavioural scores, suggesting little developmental change.

**Figure 2.**
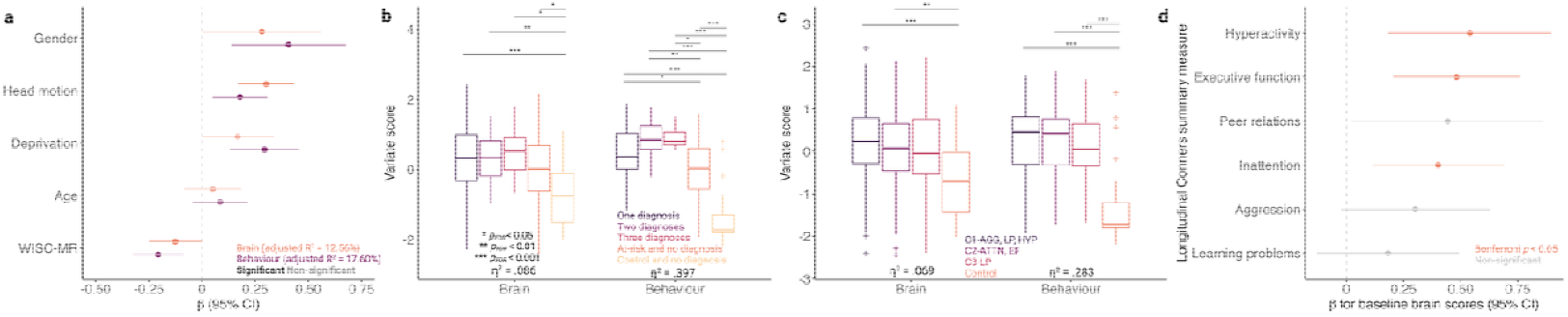
Sensitivity of brain and behavioural scores to demographics, diagnostic burden and symptom severity, and potential prognostic value for longitudinal psychopathology. **a**. Fluid reasoning was quantified by the matrix reasoning scale from the Weschler Intelligence Scale for Children (WISC-MR). Deprivation was measured by the index of multiple deprivation (IMD), reverse-coded and normalized relative to the largest rank, such that higher IMD scores represented higher deprivation. Head motion was measured as mean framewise displacement, and age as age at scan. **b**. Participants were grouped into 5 groups based on diagnostic burden: at-risk with no diagnosis (N = 109), comparison with no diagnosis (N = 26), 1 diagnosis (N = 42), 2 diagnoses (N = 16), and 3 or more diagnoses (N = 8). The most common diagnosis was combined ADHD (N = 33), followed by those undergoing speech and language therapy (N = 32), those with dyslexia (N = 17), autism (N = 13), dyspraxia (N = 7) and global developmental delay (N = 3). Rarer conditions included anxiety (N = 2), inattentive ADHD (N = 2), and a known genetic (microdeletion) condition (N = 2). Single instances were observed of dyscalculia, hyperactive ADHD, foetal alcohol spectrum disorder, depression, a diagnosed language disorder, obsessive compulsive disorder, Tourette’s, and diagnosed deficits in attention and motor perception. Of those with a diagnosis, only one participant was from the control group. Diagnoses were primarily neurodevelopmental and clinically recognised in the Diagnostic and Statistical Manual of Mental Disorders. **c**. Data-driven clustering on Conners composite scores from a prior study of 777 struggling learners in CALM (Jones et al., 2021) were used to define 3 groups in the present study. C1 (N = 74) displayed elevated aggression (AGG), learning problems (LP), and hyperactivity/impulsivity (HYP); C2 (N = 51) displayed heightened inattention (ATTN) and executive function problems (EF); whilst C3 (N = 43) displayed elevated learning problems (LP) only, relative to 27 controls. Note that data-driven groupings were missing for 6 participants in the current study, reducing the analyses to 195 participants. Across **b** and **c**, as brain scores were not normally distributed [*W*(201) = 0.988, *p* = 0.086], a Kruskal-Wallis test was used. As behavioural scores were normally distributed [*W*(201) = 0.963, *p* = 4.10 × 10^-5^ ], a one-way ANOVA was used. **d.** Baseline connectivity scores were significantly linked to longitudinal summary scores of hyperactivity/impulsivity, executive function, and inattention. Multiple comparisons were corrected for by controlling the false discovery rate (FDR). Across boxplots, the central line denotes the median, flanked by the upper and lower quartiles, tails denote the non-outlier ends of the distributions, whilst crosses denote outliers.

Next, we examined the sensitivity of brain and behavioural scores to diagnostic burden (**Figure 2b**). Diagnoses were primarily neurodevelopmental and clinically recognised in the Diagnostic and Statistical Manual of Mental Disorders (DSM). However, several conditions represented treatment for broad but otherwise undiagnosed difficulties, such as undergoing speech and language therapy. A Kruskal-Wallis test showed that higher diagnostic burden was linked to higher brain scores [*H*(4) = 20.759, *p* = 3.53 × 10^-4^, η^2^ = 0.086]. However, post-hoc comparisons were only significant between the control group with no diagnoses and all other groups, suggesting that brain scores differentiate between referral group, namely at-risk or control, but not diagnostic burden thereafter. Stronger diagnostic burden was significantly associated with higher behavioural scores [*F*(4, 196) = 32.244, *p* = 1.20 × 10^-20^, η^2^ = 0.397], with an effect almost four-times that of brain scores. Post-hoc Dunn tests with Bonferroni correction showed a greater sensitivity of behavioural scores to diagnostic burden, distinguishing not only between referral groups but also between number of diagnoses. To examine effects of symptom profile on variate scores (**Figure 2c**), we used prior data-driven transdiagnostic groupings based on the Conners scale (Jones et al., 2021). Symptom profiles exerted a significant effect on both brain [*H*(3) = 16.202, *p* = 0.001, η^2^ = 0.069] and behavioural scores [*F*(3, 191) = 25.072, *p* = 1.01 × 10^-13^, η^2^ = 0.283], differentiating between referral group only.

### Baseline connectivity scores pre-empt longitudinal psychopathology

To evaluate the extent of overfitting to baseline psychopathology, we evaluated potential associations between baseline PLS-derived connectivity scores and 6 longitudinal age-standardised Conners psychopathology composite scores, for a subset of 85 referred CALM children with longitudinal data, followed up on average 60.20 months (± 11.46) after baseline scanning. In each linear model, we controlled for gender, follow-up interval, baseline head motion and baseline deprivation. As shown in **Figure 2d**, baseline connectivity scores were significantly linked to higher age-standardised Conners scores for hyperactivity/impulsivity (β = 0.540, 95% CI: [0.183, 0.897], *p*_*FDR*_ = 0.004, *p*_FDR_ = 0.011, adjusted *R*^2^ = 18%), executive functioning (β = 0.481, 95% CI: [0.204, 0.758], *p*= 0.001, *p*_*FDR*_ = 0.005, adjusted *R*^2^ = 21%), and inattention (β = 0.402, 95% CI: [0.113, 0.690], *p* = 0.007, *p*_*FDR*_ = 0.014, adjusted *R*^2^ = 15%), alongside peer relations at an uncorrected threshold (β = 0.443, 95% CI: [0.023, 0.863], *p* = 0.039, *p*_*FDR*_ = 0.059, adjusted *R*^2^ = 8%). This suggests that baseline connectivity scores have potential longitudinal prognostic value and thus generalise beyond a single time point.

### The relationship between FC and psychopathology is spatially constrained by multi-modal cortical maps

Using spatial correspondence analyses, we tested the relationship between the predictive value of functional connectivity for psychopathology, measured through the bootstrap ratio (BSR), with seven von Economo cytoarchitectonic classes (von Economo & Koskinas, 1925), four Mesulam levels of laminar differentiation (Mesulam, 2000), seven adult normative maps of MEG frequency bands from the Human Connectome Project (Van Essen et al., 2013), and seven cell type distribution maps (Seidlitz et al., 2020). For cytoarchitectonic classes and levels of laminar differentiation (**Figure 3a**), we conducted two sets of analyses. The first examines the distribution of BSRs *across* classes in a descriptive manner, whilst the second assesses enrichment *within* classes relative to spatial permutation tests. For cytoarchitectonic classes, we observed a significant effect of class on mean BSRs [*H*(6) = 15.404, *p* = 0.017]. Specifically, FC within the primary motor (*M* = 0.09, *SD* = 0.348), first (*M* = 0.172, *SD* = 0.359) and second (*M* = 0.177, *SD* = 0.449) association classes, alongside the limbic class (*M* = 0.152, *SD* = 0.383), conferred increased risk for psychopathology, whilst FC within the primary/secondary sensory (*M* = -0.018, *SD* = 0.514), primary sensory (*M* = -0.140, *SD* = 0.482) and insula classes (*M* = -0.002, *SD* = 0.479) conferred resilience against psychopathology. A post-hoc Dunn test demonstrated that the only significant comparison was between the first association class and the primary sensory cortex (*p* = 0.047), occupying separate ends of the S-A axis, development of which has been linked to psychopathology (Sydnor et al., 2021). In the second analysis, we compared the empirical mean BSR for each partition with the equivalent derived from 5000 null distributions with preserved spatial autocorrelation. Stronger FC was linked to lower psychopathology within the primary/secondary sensory (*p*_*spin*_ = 0.011, Cohen’s *d* = 2.481) and primary sensory cortices (*p*_*spin*_ = 0.011, *d* = 2.436), suggesting a protective effect. However, the BSRs did not significantly vary across classes of laminar differentiation [*H*(3) = 5.344, *p* = 0.148], nor where they significantly enriched within classes. For MEG frequency bands and cell type distributions (**Figure 3b**), we compared the empirical correlation between each map and the BSRs, against the equivalent derived from 5000 null distributions with preserved spatial autocorrelation. The BSRs were negatively correlated with α (*r* = -0.077, *p*_*spin*_ = 0.034) but positively correlated with high-γ (*r* = 0.147, *p*_*spin*_ = 0.012) and θ (*r* = 0.037, *p*_*spin*_ = 0.047). Further, BSRs were positively correlated with the expression of cortical astrocytes (*r* = 0.153, *p*_*spin*_ = 0.012) and excitatory neurones (*r* = 0.139, *p*_*spin*_ = 0.021). Together, this suggests that the predictive value of FC for psychopathology is spatially constrained by selective divisions of cytoarchitecture, MEG frequencies, and cell type distributions.

**Figure 3.**
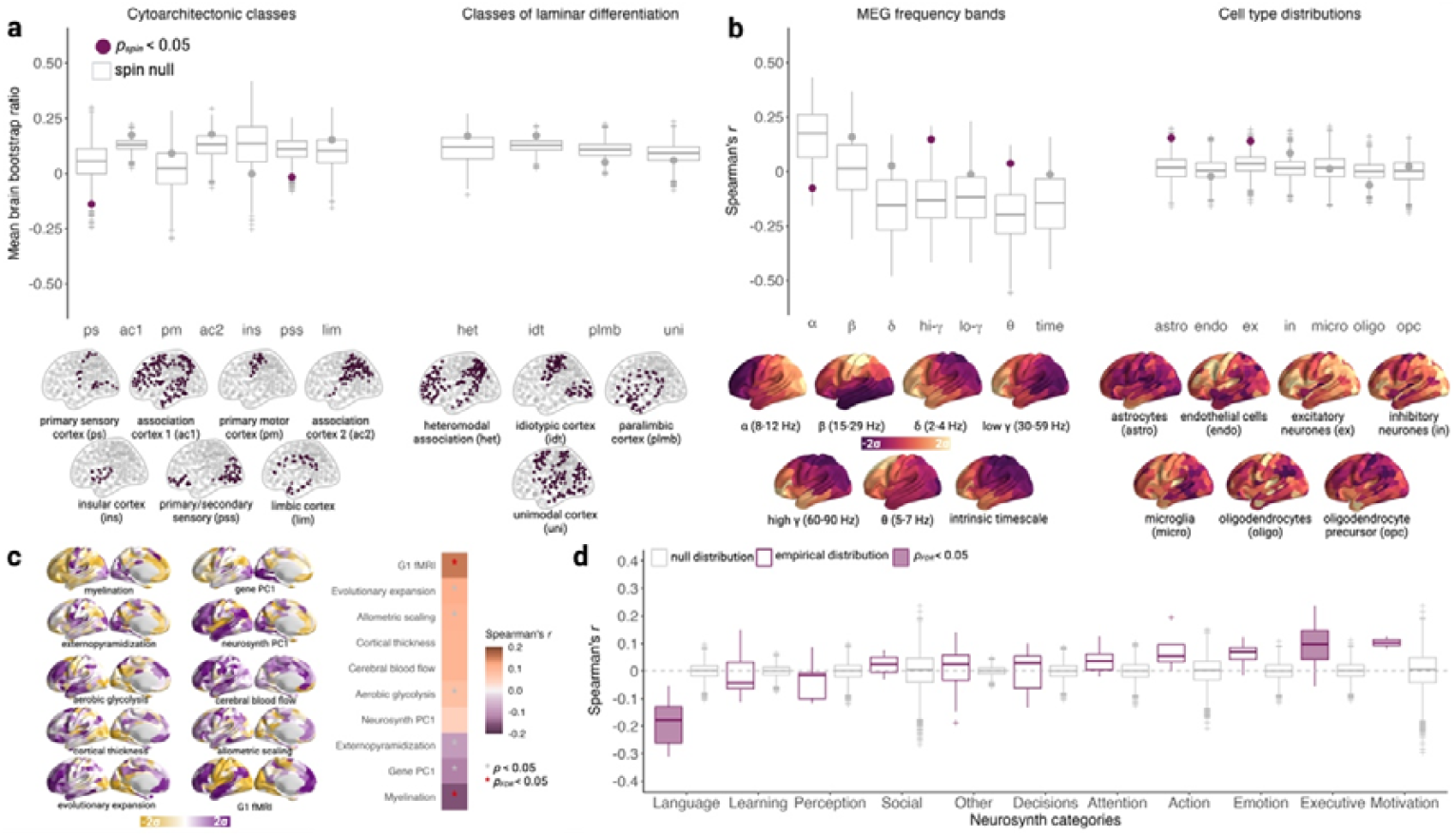
Benchmarking the relationship between functional connectivity and psychopathology against hierarchies of cortical organisation, biological maps, and cognitive terms. **a.** The mean bootstrap ratio (BSR), representing the relationship between FC and psychopathology, was calculated for 7 cytoarchitectonic classes (von Economo & Koskinas, 1925), namely primary motor (N = 26), association 1 (N = 155), association 2 (N = 77), primary/secondary sensory (N = 64), primary sensory (N = 23), limbic (N = 39) and insula (N = 16). This was repeated for 4 classes of laminar differentiation (Mesulam, 2000), namely paralimbic (N = 61), heteromodal association (N = 136), unimodal (N = 120), and idiotypic (N = 83). **b.** We obtained prior processed (Shafiei et al., 2023) canonical MEG power spectra data across 6 frequencies and intrinsic timescale, from the Human Connectome Project (Van Essen et al., 2013). Cell type distributions were obtained from prior work (Shafiei et al., 2023) in which the expression of each cell type in the Allen Human Brain Atlas microarray data was calculated using gene sets derived from cell type deconvolution (Seidlitz et al., 2020). **c.** Nodal mean BSRs were correlated with the 10 maps comprising the sensorimotor-association axis (Sydnor et al., 2021). References and brief methodological details for each map are provided in the Methods section titled ‘Alignment with the S-A axis’. **d.** We calculated the correlation between the BSR brain map and probabilistic functional associations linked to each of 123 terms from Neurosynth (Yarkoni et al., 2011). These were grouped into 11 classes: motivation (N = 2), social function (“social”, N = 2), action (N = 4), attention (N = 9), executive and cognitive control (“executive”, N = 10), perception (N = 10), emotion (N = 10), reasoning and decision-making (“decisions”, N = 11), language (N = 12), learning and memory (“learning”, N = 21), and others (N = 32). Null models (grey) were formed by randomly permuting category labels 5,000 times. Associations in **a** and **b** (two-tailed) were assessed using 5,000 null models preserving spatial autocorrelation (Váša et al., 2018) and are plotted in grey. In **a**, statistical significance was determined by the mean nodal BSR for each class after class labels were permuted using the null model. In **b**, statistical significance was determined by the magnitude of the empirical correlation coefficient compared to the null. Across all boxplots, the central line denotes the median, flanked by the lower and upper quartiles. The ends of the boxplot correspond to non-outlier ends of the distribution, whilst crosses denote outliers.

We contextualised the biological constraints associated with the relationship between FC and psychopathology by making comparisons with functional, macrostructural, microstructural, and evolutionary maps comprising the S-A axis (**Figure 3c**; Sydnor et al., 2021). In the Discussion, we explore the relative developmental stability of such maps. Of note, we highlight recent work by Monaghan and colleagues (2025), who demonstrated that the principal gradient of FC in childhood is conserved throughout adolescence and adulthood, undergoing gradual refinement of integration across development, but not replacement, and thus may be interpreted as equivalent to the adult-derived FC gradient constituting the S-A axis. The distribution of BSRs was significantly positively correlated with the principal component of adult normative FC (*r* = 0.154, *p*_*spin*_ = 0.001, *p*_*FDR*_ = 0.026), evolutionary expansion (*r* = 0.114, *p*_*spin*_ = 0.016, *p*_*FDR*_ = 0.120), allometric scaling (*r* = 0.103, *p*_*spin*_ = 0.027, *p*_*FDR*_ = 0.125), and aerobic glycolysis (*r* = 0.081, *p*_*spin*_ = 0.030, *p*_*FDR*_ = 0.125). This suggests that regions in which FC predicts higher psychopathology tend to be transmodal, with high evolutionary expansion, allometric scaling, and glucose metabolism. The distribution of BSRs was significantly negatively correlated with cortical thickness (*r* = -0.158, *p*_*spin*_ = 0.002, *p*_*FDR*_ = 0.026) and externopyramidization (*r* = -0.091, *p*_*spin*_ = 0.027, *p*_*FDR*_ = 0.125). This suggests that regions in which FC predicts lower psychopathology tend to be lightly myelinated and dominated by infragranular feedback connectivity. We further investigated whether these associations were driven by the alignment between each S-A axis composite map and the overall S-A axis rank. Using a series of partial correlations, we found that the relationships between the distribution of BSRs and each S-A composite map were non-significant after controlling for the S-A ranks overall (all *p* > 0.376). This suggests that the alignment between the distribution of BSRs and each composite map may be driven by the magnitude of the map-specific alignment with the overall S-A axis.

Recognising that the association between the psychopathology LV and the principal component of adult-derived gene expression may limit the developmental inferences we can make, we subsequently explored correspondence with three *higher-order* orthogonal AHBA genetic organisational axes (Dear et al., 2024), where the primary two components (C1 and C2) exhibit minimal developmental change, whilst the third component C3 exhibits greater developmental change by emerging through adolescence specifically. Methodological details are provided briefly in the Methods section and in the original paper. The first two genetic components are thought to be linked to autism spectrum conditions, whilst the third may be linked to schizophrenia and represents a normative transcriptional signature for cortical development specific to adolescence (Dear et al., 2024). C2 and C3 were not included in the original S-A composition: C2 was not significantly associated with the S-A axis (*r* = 0.170, *p*_*spin*_ = 0.466), whilst C3 showed moderate alignment (*r* = 0.343, *p*_*spin*_ = 0.007). We provide preliminary evidence that the distribution of BSRs was linked to C3 (*r* = 0.237, *p*_*spin*_ = 0.046, *p*_*FDR*_ = 0.069) but not C2 (*r* = 0.123, *p*_*spin*_ = 0.459, *p*_*FDR*_ = 0.459). Note that the BSRs were more strongly aligned to C1 than C3 [Steiger’s *Z* = -7.432, *p* = 3.189 × 10^-12^]. Interestingly, the relationship between C3 and the BSR distribution held even after controlling for S-A rank using a partial correlation [*r*(156) = 0.196, 95% CI = [0.04, 0.34], *p* = 0.015]. This suggests that the neurodevelopmental FC signature captured by our latent variable may generalise to overlapping risk of schizophrenia and enrichment within adolescence specifically.

### The FC-psychopathology multivariate pattern overlaps with that of language and executive functioning

Finally, to contextualise the cognitive and behavioural associations of the relationship between FC and psychopathology, we compared the distribution of BSRs with probabilistic associations between voxel activation and 123 functional keywords from the meta-analytic engine Neurosynth (Yarkoni et al., 2011), grouped into 11 themes from the Cognitive Atlas (**Figure 3d**). The mean correlation between each functional keyword activation map and the BSR map was computed for each category and compared against 5000 null distributions in which the category labels were shuffled. The BSR distribution was significantly negatively correlated with the distribution of probabilistic functional activations linked to language (mean *r* = -0.185, *p*_*FDR*_ < 0.001). This suggests that in regions most frequently activated by language tasks, stronger FC predicts lower psychopathology, introducing functional integrity of language networks as a possible protective factor against psychopathology. For context, language was most linked to functional activations within the temporal and prefrontal cortex portions of the default-mode RSN, alongside the first association cytoarchitectonic class and unimodal laminar differentiation. On the other hand, the BSR distribution was significantly positively correlated with functional activation of networks related to executive functioning (mean *r* = 0.103, *p*_*FDR*_ = 0.007). This suggests that in regions strongly activated during tasks probing executive functions, stronger FC predicts higher psychopathology, thus conferring risk. For context, executive functioning was most linked to functional activity within bilateral right hemisphere portions of the fronto-parietal RSN, alongside the second association cytoarchitectonic class and heteromodal laminar differentiation. Of note, whilst the spatial pattern of probabilistic functional associations for executive functioning (*r* = 0.200, *p* < 0.001) were significantly related to the S-A axis, language was not (*r* = 0.041, *p* = 0.418), thus highlighting language as an additional axis of variability along which the BSRs are patterned.

### Sensitivity analyses

We performed several sensitivity analyses to assess the impact of analytical choices on our results. First, to assess the effect of parcellation resolution, we repeated the PLS using the coarser Schaefer 100-node parcellation, comprising 4,950 unique edges. A single component accounted for 45.24% of FC-psychopathology covariance (permuted *p* = 0.009). Across parcellations, brain (*r* = 0.959, *p* < 0.001) and behavioural (*r* = 0.999, *p* < 0.001) scores were strongly correlated, suggesting that both methods captured similar brain-behaviour relationships irrespective of parcellation. However, the distribution of BSRs from the Schaefer 100-node parcellation were not significantly aligned with the S-A axis (*r* = -0.075, *p* = 0.458), suggesting that alignment with the S-A axis may only emerge at finer spatial resolutions. Second, to assess the effect of over-fitting in the PLS, we repeated the main analysis using a regularized canonical correlation analysis (CCA; Witten et al., 2009). CCA maximises the correlation between input matrices, rather than covariance in the case of PLS, and pre-whitens the inputs, meaning that within-set correlations are corrected for and will not drive the relationship between predictors and responses (Hansen et al., 2021). The L2-regularization term (set at 0.7) enforces sparsity and thus reduces over-fitting. Across the PLS-derived and regularized CCA-derived solutions, the brain (*r* = 0.979, *p* < 0.001) and behavioural (*r* = 0.957, *p* < 0.001) scores were highly consistent, alongside the distribution of the 79,800 brain weights (*r* = 0.962, *p* < 0.001). Third, given that head motion was correlated with brain and behavioural subject scores (**Figure 2a**), we repeated the PLS across 5 motion quantiles (top 90%, 80%, 70%, 60%, and 50%). The distribution of bootstrap ratios (1000 bootstraps) across motion quantiles were consistently strongly correlated to that of the full sample (*r* = 0.941, *r* = 0.899, *r* = 0.864, *r* = 0.838, *r* = 0.749, all *p* < 0.001). To further guard against motion, we repeated the PLS analysis with residuals after regressing head motion from imaging and psychopathology data: the consequent bootstrap ratios were significantly strongly correlated with that of the original analysis (*r* = 0.977, *p* < 0.001). Together, this suggests that the predictive value of FC associated with psychopathology did not rely on motion.

## Discussion

Using a transdiagnostic approach, we benchmarked the multivariate relationship between FC and neurodevelopmental psychopathology, as a latent variable, across multiple levels of brain organisation. In this way, we attempted to build a multi-layered framework - spanning biological maps, brain connectivity, and behaviour - to begin to address the need for integrated neuroscientific analyses proposed by RDoC (Insel et al., 2010). Crucially, and in contrast to both group-level case-control designs and community samples, the extensive comorbidity, phenotypic complexity, and oversampling of children at risk of psychiatric and neurodevelopmental conditions, but otherwise undiagnosed, in CALM reflects that of everyday practice for psychologists and psychiatrists. Participant-level expression of the latent variable was sensitive to the presence of a diagnosis but did not differentiate between children with a confirmed diagnosis and those *at risk* of neurodevelopmental conditions, casting doubt on the utility of diagnosis alone as a neurobiologically-informed categorisation system. The relationship between FC and neurodevelopmental psychopathology was differentially constrained by different levels of cortical organisation, demonstrating a non-redundancy (Hansen et al., 2023): it followed a sensorimotor-association axis, was selectively enriched within visual, dorsal attention and fronto-parietal intrinsic functional resting state networks, but otherwise unconstrained by laminar differentiation. The latent variable was significantly spatially aligned to the distribution of cortical astrocytes and excitatory neurones, as well as select MEG canonical frequency bands. Further, using the meta-analytic Neurosynth database, we highlighted functional connectivity related to language as a potential protective factor against psychopathology, and functional connectivity related to executive functioning as a potential risk factor.

We build upon prior literature which has demonstrated that the transdiagnostic relationship between functional connectivity and psychopathology is constrained by archetypal axes of cortical organisation, such as the sensorimotor-association axis (Royer et al., 2024), as well as intrinsic functional connectivity networks (Kebets et al., 2019; Xia et al., 2018; Xiao et al., 2024). However, by triangulating overlapping levels of cortical organisation, we hoped to piece together which cortical partitions are most susceptible towards expressing particularly strong positive or negative relationships between connectivity and psychopathology. For example, we observed positive enrichment of the latent variable across the lower-frequency MEG θ band and negative enrichment within the higher-frequency α band, possibly reflecting different modes of neuronal communication. Recent work has proposed that θ-α communication represents a frequency-specific loop of information flow, with a posterior-to-anterior pattern of information flow in the α band between parieto-occipital and frontal regions, and the opposite pattern of flow in the θ band (Hillebrand et al., 2016). This suggests that the relationship between functional connectivity and psychopathology aligns with frequency-dependent modes of cortical organisation. Further, the θ-α enrichments may represent the generation of underlying functional connectivity and, by association, psychopathology: similar phase-amplitude coupling between lower-frequency and higher-frequency MEG bands, such as α and θ, segregate into networks with a similar topology to intrinsic functional connectivity resting-state networks (reviewed by Baillet, 2017; Florin & Baillet, 2015).

Whilst neurodevelopmental diagnosis holds some merit as a classification system, such as facilitating communication with parents and highlighting individuals who may benefit from medication (Thapar et al., 2017), the current study urges caution towards an over-reliance on diagnosis. Namely, in contrast to a categorical case-control approach to developmental psychopathology, based on diagnosis, the current study explored the relationship between functional connectivity and transdiagnostic continuous dimensions of neurodevelopmental psychopathology, akin to a *p* factor (Caspi et al., 2014), across a phenotypically diverse sample. Interestingly, we demonstrated that the participant-level expression of the connectivity signature of the latent variable was sensitive to the presence of a diagnosis, irrespective of diagnostic load, but was unable to differentiate between those with a confirmed diagnosis and those at elevated risk of neurodevelopmental conditions. The poor correspondence of connectivity scores to diagnostic burden suggests that diagnostic status holds less significance than diagnostic *vulnerability* and calls into question conclusions drawn from traditional case-control designs in which at-risk children may have otherwise been assigned to a control group due to their lack of diagnosis. Of note, we observed a significant effect of symptom category on the contribution of each behavioural item to the latent variable, whereby items related to inattention were most closely aligned to the connectivity signature, and learning problems the least.

The relationship between FC and psychopathology was spatially patterned along the sensorimotor-association axis. This suggests that aspects of individual differences underlying brain connectivity and psychopathology may be captured by a basic principle of organisation derived from macroscale, microscale, metabolic, and evolutionary maps (Sydnor et al., 2021). This is in line with prior work within a community-based sample of almost 10,000 preadolescents which demonstrated alignment within a neurodevelopmental dimension of psychopathology, and even stronger alignment within a general p-factor dimension of psychopathology (Royer et al., 2024). However, whilst tempting to conclude *definitively* that individual differences in brain connectivity and psychopathology may be reduced to a single lower-order manifold, we urge caution. Alignment with the S-A axis was weak and only emerged at a finer spatial resolution (see *Sensitivity Analyses*). First, whilst the S-A axis is a dominant gradient of cortical organisation, and benefits from a well-developed theoretical framework (Sydnor et al., 2021), it is not the only one. For example, an intrinsic two-dimensional coordinate system has been recently proposed, comprising of a sensorimotor-transmodal primary axis, and a secondary axis differentiating between visual regions and those linked to somatosensory and auditory processing (reviewed by Huntenburg et al., 2018). Further, we provided preliminary evidence that the spatial patterning of the latent variable was significantly linked to higher-order dimensions of cortical gene expression only weakly linked to the sensorimotor-association axis, thus incorporating additional unique axes of organisation. Second, we find that the FC-psychopathology relationship is more strongly spatially aligned with some maps comprising the S-A axis, such as myelination and the principal component of functional connectivity, than others. The specificity of these relationships highlights the *non-redundancy* of different levels of organisation captured by the S-A axis, and their relevance to the relationship between FC and psychopathology. However, it remains unclear whether each biological map is indeed a stronger predictor of psychopathology, or whether it is simply more strongly linked to the functional correlates of that psychopathology.

We also acknowledge that the developmental stability of each composite map is modality-dependent. For example, the principal component of functional connectivity recapitulates a unimodal-transmodal axis by 6 years of age, and persists throughout adolescence and adulthood (Monaghan et al., 2025), rather than being replaced by a secondary gradient at 12 years of age as previously suggested (Dong et al., 2021). This implies that whilst benchmarking our psychopathology latent variable against a child-derived FC gradient may yield stronger alignment compared to its adult-derived counterpart, such increases would be minimal. However, cortical gene expression may be one modality in which developmental change is more evident. The BrainSpan atlas provides cortical and subcortical gene expression estimates across 16 unique regions from 42 donors ranging in age between 8 post-conceptional weeks to 40 years old (Miller et al., 2014), thus at a considerably sparser spatial resolution but expanded age range compared to the AHBA-derived components used in the present study. Dear and colleagues (2024) demonstrated that the principal two components of developmental gene expression derived from the BrainSpan atlas across pre-birth and the period between birth and 13 years old were consistent with corresponding AHBA components, suggesting that the principal two axes of gene expression are at an adult-form early in life (all *r* ≥ 0.66). Yet, such correlations are moderate, suggesting that some developmental finetuning occurs, particularly in the case of the third component whose AHBA-BrainSpan correlation was weak during pre-birth (*r* = 0.29) and increased across development. Together, this suggests the utility of such gradients as theoretical scaffolds upon which developmental trends unfold depends on their modality-specific stability. In more broad terms, this also implies that our understanding of developmental stability of cortical axes is ever-evolving.

In examining the relationship between brain connectivity and psychopathology, there exists a continuum of approaches. One end of the continuum is anchored by focused and discrete subtyping, where multiple orthogonal components are derived linking brain connectivity and psychopathology. This allows us to explore within-diagnosis heterogeneity, typically within a narrower symptom space. For example, differential expression of ASD-related genes, particularly those related to immune signalling, may explain individual differences in atypical functional connectivity of ASD subtypes across verbal intelligence, social affect, and restricted and repetitive behaviours (Buch et al., 2023). The current study sits at the opposite end of this continuum, creating a continuous measure of variability shared by all participants. Both approaches allow for modelling of a diversity of neurodevelopmental pathways towards the same phenotypic outcome, namely equifinality (Cicchetti & Rogosch, 1996), but differ in the types of variability, and therefore biological associations, they capture.

We outline three possible limitations and extensions of our work. The first relates to the generalizability of neurodevelopmental psychopathology to general psychopathology. Within a hierarchical transdiagnostic framework, neurodevelopmental psychopathology emanates from broad externalizing difficulties, which in turn are a specialised sub-spectrum of general psychopathology (Caspi et al., 2014; Holmes et al., 2019; Kotov et al., 2017). A recent study has suggested partial overlap between the two, whereby the relationship between functional connectivity and psychopathology are patterned across an S-A axis in both general and neurodevelopmental psychopathology (Royer et al., 2024). However, multimodal prediction of the *p* factor and the neurodevelopmental dimension also demonstrate important differences in sensitivity by region and modality. These include widespread reductions in cortical volume, thickness, fractional anisotropy and mean diffusivity across all tracts in general psychopathology, compared to more localised increases in surface area and volume within temporoparietal regions, accompanied by decreased fractional anisotropy within portions of the cingulate cingulum and longitudinal fasciculus specific to neurodevelopmental psychopathology (Royer et al., 2024). The onset of neurodevelopmental conditions, typically in childhood, predates the onset of mood- and anxiety-based conditions, typically during adolescence (Solmi et al., 2022). Therefore, it is unclear to what extent the biological basis of neurodevelopmental conditions *generalizes* to that of broader psychopathology. This contributes to a wider debate concerning whether biological and connectivity correlates for disorders overall are stable across time or whether they emerge as psychopathology later in life due to environmental stress. In the current study, we find that the connectome variate scores are stable across development, supporting the former hypothesis. Further work deriving orthogonal components of psychopathology – mood, psychosis, externalizing behaviour, and fear – have shown that whilst the connectivity basis of some dimensions display developmental increases, specifically for mood and psychosis, the connectome signatures of externalizing behaviours, onto which ADHD most reflects, may be stable across time (Xia et al., 2018). Another study has begun to unravel the overlap between general psychopathology and neurodevelopment (Vedechkina et al., 2025). Namely, overall genetic liability for general psychopathology may be distinct to that of neurodevelopmental conditions, with the latter conferring possible resilience against adversity during pre-adolescence (Vedechkina et al., 2025). Ultimately, pooling together phenotypically-rich longitudinal data across the entirety of childhood and adolescence and examining the connectivity signatures of pre-defined taxonomic hierarchies of psychopathology spanning both general and neurodevelopmental dimensions, will be the first step towards disentangling the temporal relationship between general psychopathology and its sub-strata.

A second possible limitation and extension of our work focuses on our use of functional connectivity. Structural connectivity, consisting of physical white matter axonal projects connecting pairs of regions, provide a sparse structural backbone from which dynamic functional connectivity emerges (Honey et al., 2009). Most functional connections do not have direct structural support, thus emerging as higher-order interactions, with structure-function correspondence following a unimodal-transmodal hierarchy, with tight coupling at the unimodal pole (see Suárez et al., 2020). Multivariate techniques are poised to relate variation between predictors and outcomes. The increased variability of function, compared to structure, thus is an appealing imaging measure to relate to psychopathology, and one which has been extensively examined (Royer et al., 2024; Xia et al., 2018; Xiao et al., 2024). However, multimodal neuroimaging studies have demonstrated non-redundancy between metrics within and across neuroimaging modalities (Jirsaraie et al., 2025; Liu et al., 2023; Royer et al., 2024; Tsvetanov et al., 2025). Specifically, a recent study systematically compared the predictive power of white matter, grey matter, and functional connectivity attributes for a tri-factor psychopathology model. They demonstrated that multimodal models yielded the greatest predictive accuracy, followed by white-matter neurite orientation dispersion and density imaging, cortical thickness, and only then functional connectivity (Jirsaraie et al., 2025). The superior performance of multimodal models suggests that both structural and functional measures provide non-redundant insights into the aetiology of psychopathology. One way in which structural MRI features may be integrated with functional connectivity, in a low-dimensional manner, within a predictive framework for psychopathology is through morphometric similarity networks (MSNs; Seidlitz et al., 2018). For each participant and node, MSNs are constructed by correlating multiple structural MRI measures, providing a more comprehensive insight into structural form.

A third limitation of our work reflects the high dimensionality of the underlying latent variable compared to the modest sample size, particularly pertinent in the case of relatively low split-half reliability. Whilst sample sizes into the thousands are ideal, including those afforded by UK Biobank and the Human Connectome Project as typically-developing cohorts, and yield highly stable brain-behaviour relationships (Helmer et al., 2024), such requirements are less attainable with clinical developmental cohorts, like CALM, at least at present. Safeguarding against overfitting may be facilitated in other ways, such as through regularization in sparse CCA, external validation, enrichment of biological maps, and evaluating longitudinal prognostic value of brain-behaviour relationships. Whilst large consortia are one pathway towards establishing reliable brain-behaviour relationships, targeted investigation of smaller clinical groups with a greater signal-to-noise ratio than their typically-developing counterparts are another route (Gratton et al., 2022).

In summary, we derived a temporally-stable transdiagnostic functional connectivity signature of neurodevelopmental psychopathology, spanning childhood and adolescence. We benchmarked this signature across multiple cortical maps, building an analysis framework spanning biological maps, cortical connectivity, and psychopathology. In this way, we have begun to reconcile changes in cortical connectivity related to psychiatric and neurodevelopmental conditions with their underlying biological underpinnings, to develop more reliable and biologically-informed biomarkers of childhood psychopathology. This signature was circumscribed by anatomical and functional cortical organisational hierarchies, was spatially aligned with canonical MEG frequency bands alongside distributions of cortical astrocytes and excitatory neurones and displayed significant overlap with functional networks activated by language and executive functioning. Further, the connectivity signature was significantly, albeit weakly, aligned to an archetypal sensorimotor-association axis, with connectivity predicting greater psychopathology at the associative end. Our findings call into question the overuse of diagnostic labels in psychiatric research, and we situate our findings within the wider context of the relationship between general psychopathology and its substrata, as well as predicting behaviour using neuroimaging.

